# Control of septin filament flexibility and bundling

**DOI:** 10.1101/209643

**Authors:** Anum Khan, Jay Newby, Amy S. Gladfelter

## Abstract

Septins self-assemble into heteromeric rods and filaments to act as scaffolds and modulate membrane properties. How cells tune the biophysical properties of septin filaments to control filament flexibility and length, and in turn the size, shape, and position of higher-order septin structures is not well understood. We examined how rod composition and nucleotide availability influence physical properties of septins such as annealing, fragmentation, bundling and bending. We found that septin complexes have symmetric termini, even when both Shs1 and Cdc11 are coexpressed. The relative proportion of Cdc11/Shs1 septin complexes controls the biophysical properties of filaments and influences the rate of annealing, fragmentation, and filament flexibility. Additionally, the presence and exchange of guanine nucleotide also alters filament length and bundling. An Shs1 mutant that is predicted to alter nucleotide hydrolysis has altered filament length and dynamics in cells and impacts cell morphogenesis. These data show that modulating filament properties through rod composition and nucleotide binding can control formation of septin assemblies that have distinct physical properties and functions.

## Introduction

Septins are filament-forming proteins that belong to a family of P-loop GTPases and are conserved from yeast to humans. A key feature of the septin cytoskeleton is that filaments form from multimeric complexes that show plasticity in composition depending on cell type or developmental stage. In yeast, four septins form an octameric complex where the terminal position can be occupied by two different septins: Cdc11 and Shs1 (Garcia *et al.*, 2011). Multiple distinct septin complexes also coexist in a mammalian cell with different subunit combinations and splice variants possible. (Sellin *et al.*, 2011). What is the functional consequence of this compositional heterogeneity on septin form and function?

In principal, varying the mixtures of heteromeric complexes could change the material properties of filaments, the subcellular localization and/or interaction partners. Mixing different proportions of Shs1 and Cdc11 complexes can give rise to different ultra-structures *in vitro* (Garcia *et al.*, 2011). Different septin heteromers also generate diverse higher order structures in *Aspergillus nidulans* (Hernández-Rodríguez *et al.*, 2014). Similarly, SEPT9 presence in complexes promotes different higher order structures that function at different times and places in mammalian cells (Kim *et al.*, 2011). Despite clear evidence of functional differences, little is known about how septin complex composition gives rise to these differences.

Recent evidence shows that the septin subunit order and oligomer composition is controlled by GTP binding and hydrolysis rates (Weems and McMurray, 2017). Other studies suggest GTP binding/hydrolysis may also be important for the formation of higher order septin structures both *in vitro* and *in vivo* (Versele *et al.*, 2004; Huang *et al.*, 2006; Nagaraj *et al.*, 2008; Weems *et al.*, 2014; Akhmetova *et al.*, 2015). However, there is little evidence of nucleotide hydrolysis by the septin complex over one yeast cell cycle, raising the issue of whether GTP plays a role beyond a structural one in controlling filament dynamics (Vrabioiu *et al.*, 2004). We examine how septin complex composition and nucleotide state affects filament properties to gain insight into how cells may produce structurally and functionally distinct higher-order septin structures.

## Results and Discussion

### Analysis of septin complex formation with co-expressed terminal subunits

In *Saccharomyces cerevisiae*, two septin subunits, Cdc11 and Shs1 can occupy the terminal position in a hetero-octamer. The relative abundance of complexes with different terminal septins changes the size and shape of septin ultra-structures *in vitro* (Garcia *et al.*, 2011). The overall Cdc11:Shs1 ratio is ~1:1 in yeast, although the composition and relative abundance of septin complexes has not been directly measured (Chong *et al.*, 2015). In fact, there are many possible combinations of complexes (Figure 1A).

**Figure 1.**
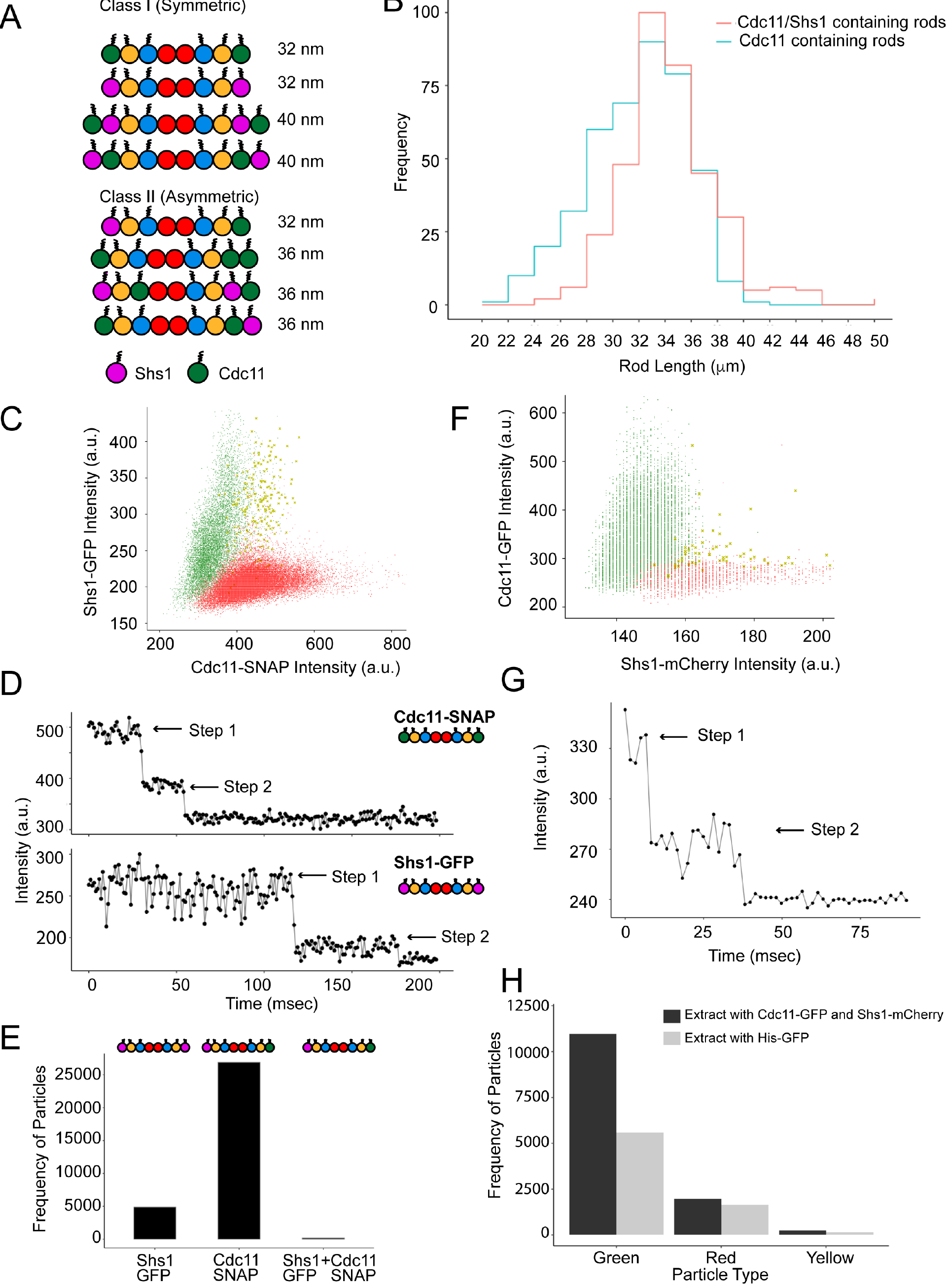
Symmetric rod composition *in vitro* and *in vivo*. (A) Schematic depiction of possible rod combinations divided into two symmetry-based classes (B) Rod length distributions from negative stain TEM images (n ≥ 400 rods) of septins in 300 mM salt. (C) Average intensity distribution for single particles over time using complexes formed when Cdc11 and Shs1 are coexpressed recombinantly. (D) Two-step photo-bleaching profiles of Cdc11-SNAP and Shs1-GFP particles. (E) Frequency of homotypic and heterotypic particles among purified septin complexes. (F) Average intensity distributions for particles pulled down by a-GFP from yeast extract containing Cdc11-GFP and Shs1-mCherry. (G) A two-step photo-bleaching profile of particles in F. (H) Frequency of green (Cdc11-GFP), red (Shs1-mCherry), and yellow (Cdc11-GFP + Shs1-mCherry) particles for extract containing Cdc11-GFP and Shs1-mCherry compared to His-GFP extract as a negative control.

We first investigated what complexes are built when Cdc11 and Shs1 are coexpressed. To measure the size distribution of complexes, we coexpressed four (Cdc10, Cdc3, Cdc12 and Cdc11) or five (Cdc10, Cdc3, Cdc12, Cdc11 and Shs1) septins, purified from *E. coli* and visualized via negative stain transmission electron microscopy (TEM). Longer complexes were observed when both Cdc11 and Shs1 were expressed as opposed to Cdc11 only (p<0.005, KS test). However, the mode for both distributions remained at 32 nm, the length of an octamer (Figure 1B). Thus, although longer complexes can form, the majority of rods are octameric when Shs1 and Cdc11 are coexpressed.

To determine whether octamers are symmetric or asymmetric (Figure 1A, class 1 v. class 2), we analyzed single septin complexes via TIRF microscopy. Purified complexes containing Cdc11-SNAP and Shs1-GFP were added at very low concentrations (pM) to detect single particles. Signal intensity analysis revealed two distinct complex populations containing either Shs1-GFP or Cdc11-SNAP, but none with both signals (Figure 1C). Two-step bleaching profiles confirmed that each complex contained two Cdc11-SNAP or two Shs1-GFP molecules (Figure 1D). A very small population of particles contained both Shs1-GFP and Cdc11-SNAP signal that was difficult to distinguish from noise (Figure 1, C and E). These data suggest that when Cdc11 and Shs1 are coexpressed, septin rods form two distinct, symmetric complexes with either Cdc11 or Shs1 present.

To determine the composition of native yeast complexes, we used single molecule pulldown (SiMPull) on extracts from cells expressing both Cdc11-GFP and Shs1-mCherry (Jain *et al.*, 2009). Anti-GFP antibodies were immobilized on a coverslip and captured puncta were analyzed for mCherry signal. Shs1-mCherry signals were not detected at the same positions as Cdc11-GFP signals suggesting that two distinct populations exist in yeast cytoplasm containing either Cdc11 or Shs1 (Figure 1F). Bleach step profiles revealed two Cdc11-GFP molecules per punctum consistent with rods containing Cdc11 on both ends (Figure 1G). Puncta detectable in the red channel did not colocalize with Cdc11 signal, and were indistinguishable from noise when compared with extract containing unlabeled septins and monomeric eGFP (Figure 1H). Therefore, yeast septins form exclusively symmetric complexes both *in vitro* and *in vivo* even in presence of alternate subunits.

Symmetric rod formation suggests a mechanism to coordinate the complex composition across tens of nanometers. How heteromeric complexes of these dimensions assemble so that the terminal positions “match” is challenging to envision. One explanation would be that subunits could be locally translated with co-translational complex assembly. Evidence supports coordinated septin translation in *Ustilago maydis*, where septin mRNA is transported and translated on endosomes (Baumann *et al.*, 2014; Zander *et al.*, 2016), and is supported by heterokaryon experiments in yeas (Weems *et al.*, 2014). Alternatively, allostery or some type of assembly selection may bias complex composition. Cdc3/Cdc12 nucleotide state may bias its incorporation with Cdc10 core dimers to promote symmetry, or the Cdc12 nucleotide state conformation can propagate through the oligomer to coordinate the termini states prior to Cdc11 or Shs1 binding.

### Shs1 changes the biophysical properties of septin filaments

What is the consequence of having two pools of septin complexes with different termini? Genetic and FRET-based assays have shown that Shs1 complexes do not polymerize with other Shs1 complexes (Finnigan *et al.*, 2015; Booth and Thorner, 2016). Additionally, changing the ratio of Cdc11:Shs1 complexes changes higher-order structure size *in vitro* (Garcia *et al.*, 2011). We investigated how the ratio of Cdc11- to Shs1-containing rods affects biophysical properties of septin filaments such as filament length, flexibility, annealing and fragmentation events, and bundling.

On supported lipid bilayers (SLBs), filament length was inversely related to Shs1-complex abundance (Figure 2A, Supplemental video 1 and 2). High Shs1 concentrations led to more dynamic filaments with increased annealing and fragmentation event frequencies (Figure 2, B and C). The septin filament persistence length also decreased (i.e. increased flexibility) with increasing proportions of Shs1 rods (Figure 2E). The flexural rigidity differences may reflect affinity differences between Shs1-Cdc11 and Cdc11-Cdc11 interactions (Booth *et al.*, 2015). Previous studies have shown that Shs1-containing complexes can associate laterally, whereas Cdc11-Cdc11 interactions are end on (Garcia *et al.*, 2011; Bridges *et al.*, 2014). In our experiments, bundling/unbundling events increased with Shs1 proportion (Figure 2D). The orientation or strength of Shs1-Cdc11 interactions may promote filament flexibility compared to end-on Cdc11-Cdc11 interactions (Kaplan *et al.*, 2015).

**Figure 2.**
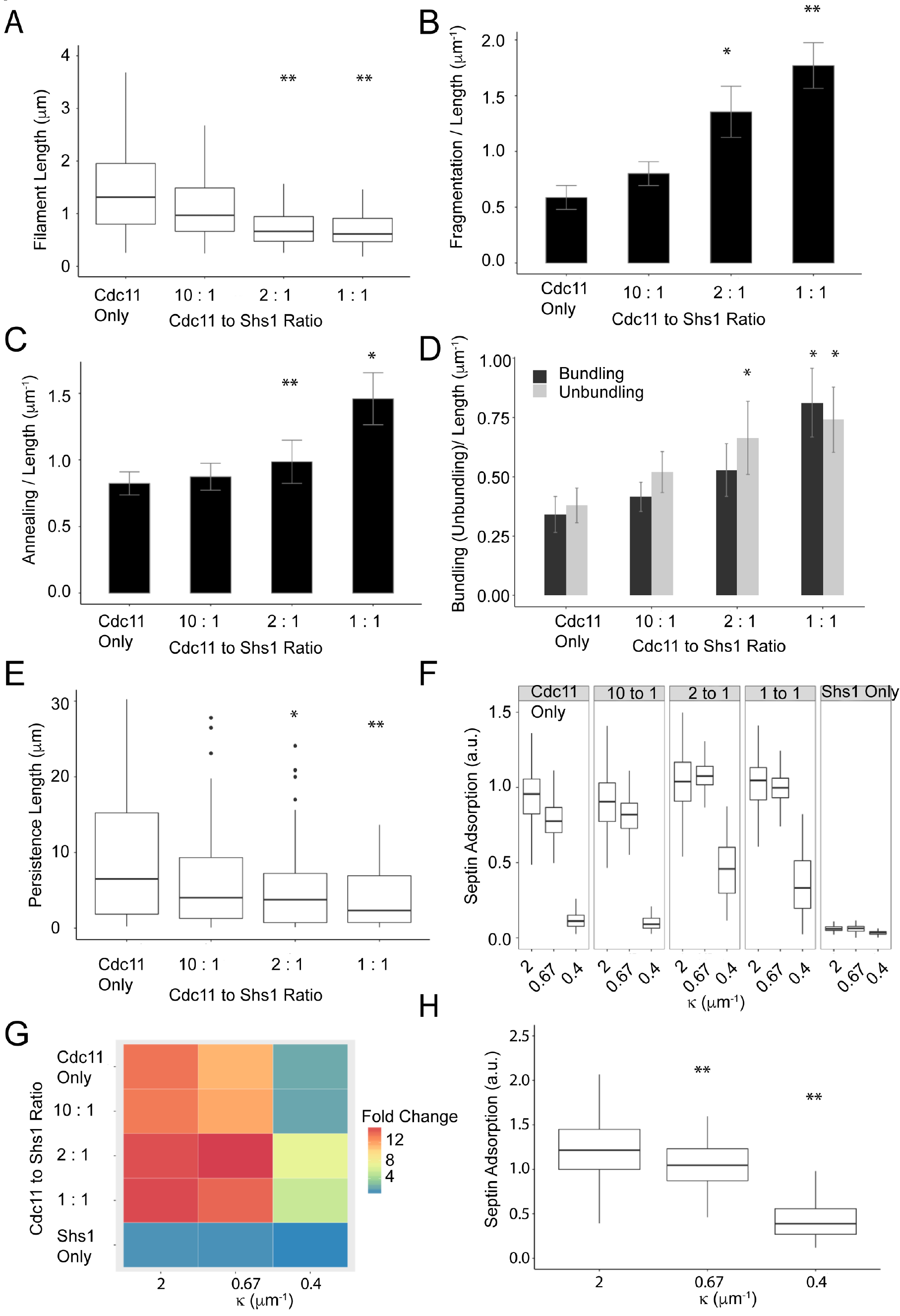
Ratio of Cdc11 and Shs1 complexes modulates septin filament properties. (A) Septin filament length (B) Fragmentation/Length (C) Annealing events/Length and (D) Bundling and unbundling per unit length with varying Cdc11:Shs1 ratios. In A, n > 250 filaments and B-D, n ≥ 49 filaments for each ratio, error bars represent standard error. *p<0.05 **p<0.005, Kolmogorov-Smirnov (KS) test. (E) Filament persistence length decreases from a median of 6.4 μm for Cdc11 only to 2.4 μm for 1:1 Cdc11:Shs1 ratio filaments (n ≥ 15 filaments). (F) Septin adsorption on different bead sizes at different Cdc11:Shs1 ratios using a total of 100 nM protein (n ≥ 50, p < 0.005 Dunn test). (G) Fold difference heat map of septin adsorption to different bead sizes with different Cdc11:Shs1 ratios, normalized to the lowest septin adsorption. (H) Shs1 complexes show curvature preference when recruited via NTA-Ni lipids using the His-tag (p < 0.005, Dunn test).

Next we found that Shs1:Cdc11 ratios do not alter the micron-scale curvature preference of septins (Figure 2F) (Bridges *et al.*, 2016). Notably, however, increasing Shs1 abundance increased total septin adsorption on beads of all sizes tested (Figure 2, F and G). This increase may arise due to differential filament packing on a curved surface, facilitated by Shs1. Shs1 complexes alone showed minimal binding at all curvatures (Figure 2, F and G), consistent with previous work showing polymerization is required for stable association to curved surfaces (Finnigan *et al.*, 2015; Booth and Thorner, 2016; Bridges *et al.*, 2016). Shs1 complexes did display a curvature preference when recruited artificially via the His-tag and Ni^2+^-NTA modified lipids (Figure 2H). Thus, Shs1 influences the flexibility, assembly, disassembly, and bundling of filaments while not changing micron-scale curvature preference.

These results suggest that varying septin complex proportions is a mechanism to tune septin biophysical properties. While Shs1 and Cdc11 are expressed approximately stoichiometrically in yeast cells, locally controlling Cdc11 and Shs1 complex ratios within the septin structure can be used to generate filaments with distinct properties and diversity in structures.

### Magnesium and guanine nucleotides promote septin filament bundles

A second prominent feature of septins that could influence filament properties is the presence and state of the guanine nucleotide. Septin complexes purified from *E. coli*, yeast, or mammalian cells contain a mixture of GTP and GDP (Field, 1996; Vrabioiu *et al.*, 2004). Recent work illuminated a key role of nucleotide state for directing the assembly order of yeast septin complexes (Weems and McMurray, 2017). It is unclear if nucleotide exchange/hydrolysis influences filament properties that arise *after* complex formation. We assessed polymerization of Cdc11-containing complexes in the presence of nucleotide and magnesium to stabilize bound nucleotide (Sirajuddin *et al.*, 2009). Strikingly, Mg^2+^ alone causes septin filaments to bundle at physiological ion concentrations (1-3 mM) both in the presence or absence of an anionic lipid (75% PC, 25% PI) SLB (Figure 3, A and B). Intensity analysis of filaments formed in presence of Mg^2+^ on SLBs shows they are twice as bright as single filaments suggesting filament pairing. We suspect that similar to F-actin, Mg^2+^ promotes bundling by shielding negative charge and reducing repulsion between actin filaments (Tang and Janmey, 1996). FtsZ studies have shown that metal ions (Mg^2+^ or Ca^2+^) bind FtsZ, displace bound GTP to a more surface exposed position, and provide a new interface for filament bundling (Marrington *et al.*, 2004). While the nucleotide binding pocket of septins is solvent inaccessible, Mg^2+^ binding may promote conformational changes of the nucleotide and the filament, promoting lateral interactions.

**Figure 3.**
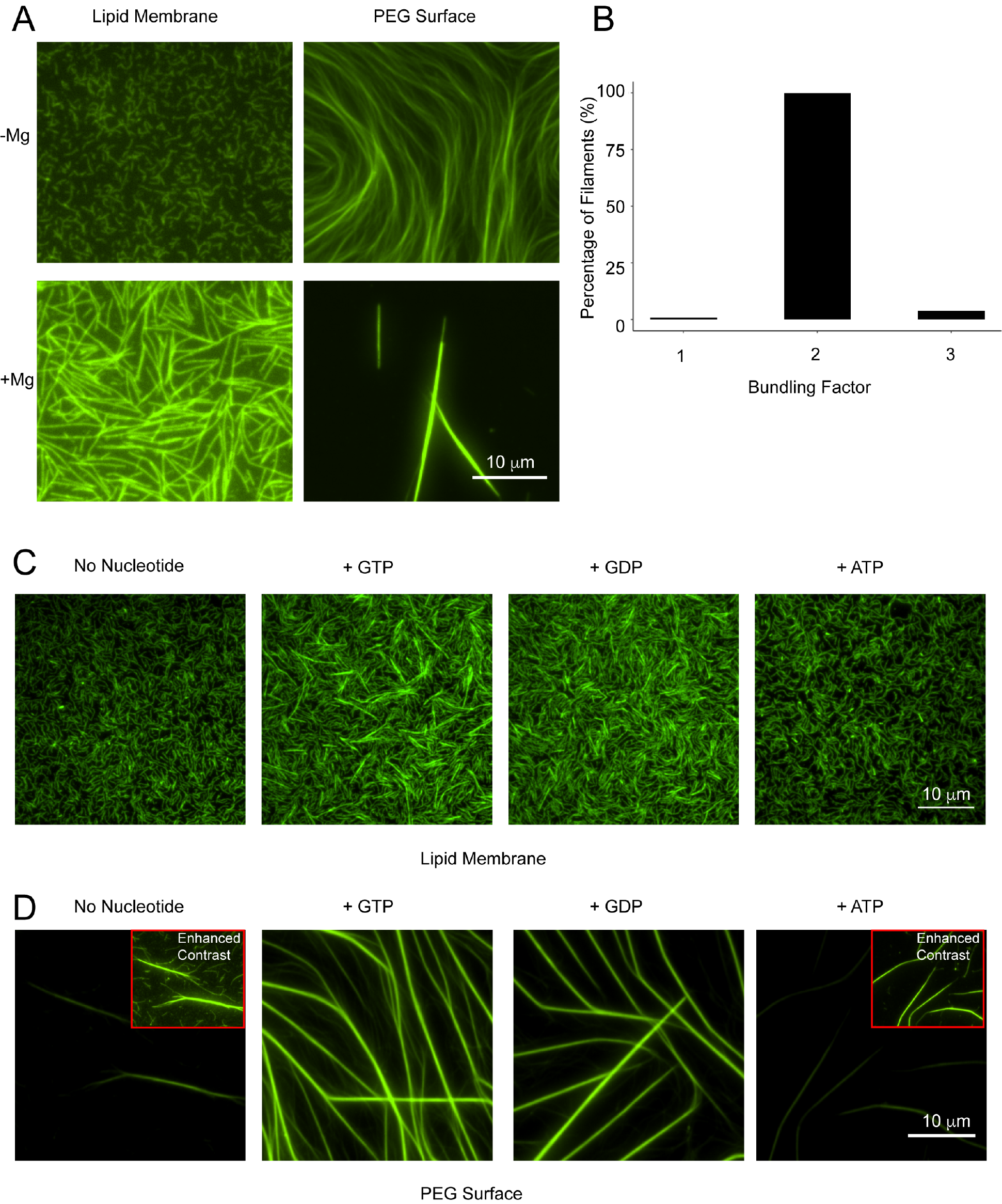
Mg^2+^ and GTP/GDP promote septin filament bundling. (A) Mg^2+^ causes septin filament bundling on SLBs and PEG. 5 nM septins were allowed to polymerize in 50 mM salt with (1 mM) and without Mg^2+^ on a SLB. 100 nM septins were polymerized in 50 mM salt on a PEG surface with (3 mM) and without Mg^2+^. (B) Filaments formed in presence of Mg^2+^ on SLBs were 2X brighter than filaments formed in absence of Mg^2+^. (A) Septins show bundling with GTP and GDP on SLBs. (D) Septins form long stable filaments in presence of GTP and GDP on PEG coated coverslips. Insets show enhance contrasted filaments. Septins were pre-incubated with excess nucleotide for 3 hours at 30°C before allowing polymerization in C and D.

To disentangle the bundling effect of Mg^2+^ from nucleotide state, we modified nucleotide exchange protocols for septins (Sheffield *et al.*, 2003; Huang *et al.*, 2006). Septins were first incubated with excess nucleotide in presence of 1 mM Mg^2+^ before diluting to 20 μM Mg^2+^ and low salt to promote polymerization (see materials and methods for details). Under these conditions, we observed normal, flexible filaments without additional nucleotide but substantial bundling in the presence of GTP or GDP. Notably, ATP did not promote bundling (Figure 3C). No differences were observed between GTP and GDP, suggesting nucleotide binding, rather than hydrolysis, promotes lateral interactions. As no bundles were detected without pre-treatment to promote exchange, increased bundling is likely due to incorporation or exchange of previously-bound nucleotide. Bundling was non-uniform throughout the membrane, possibly due to slow nucleotide turnover and Mg^2+^ requirement for GTP binding. These data suggest that nucleotide state changes can produce stiff filaments with increased lateral associations (Supplementary video 3).

Guanine-nucleotide induced bundles were longer, thicker, and sequestered more protein when septins were polymerized in solution and visualized on polyethylene glycol (PEG) treated glass instead of assembling on a SLB (Fig. 3D). This raises the possibility of a common interface between membrane binding and filament-filament interactions, indicating possible competition between these two types of interactions. Some cells normally have septin bundles, including nuclear-associated septins in mammalian cells, assemblies in yeast mating projections, and bars that specify septation in filamentous fungi (Ford and Pringle, 1991; Alvarez-Tabarés and Pérez-Martín, 2010). Cells may regulate bundle distribution relative to other types of assemblies. Previous work has also implicated the G-domain in septin filament lateral interactions. Cells lacking Cdc12 and Cdc3 coiled-coils or lacking Cdc10 and expressing a Cdc3 mutant (G261V) can generate substantial septin bundles potentially through the G-domain (Bertin *et al.*, 2010). What surfaces can promote septin filament bundling and how nucleotide state and Mg^2+^ influence bundling propensity require further study.

### Nucleotide pocket mutants show increased lateral interactions

We further examined the role of nucleotide binding in septin filament assembly by generating several mutants predicted to alter nucleotide state. A hydrolysis-defective Cdc12 mutant (T75A) and a putative gain-of-function Cdc11 mutant (D65T) were generated based on structural studies and sequence homology (Sirajuddin *et al.*, 2009). The mutants showed no complex formation defects (Figure 4A). Two other nucleotide pocket mutants (Cdc11 R35E and Cdc12 G247E) showed complex formation defects and did not form filaments (unpublished data). Both the Cdc11 D65T and Cdc12 T75A complexes formed small, bundled filaments in addition to the unbundled filaments typical of yeast septins on SLBs. The frequency of bundles was higher in Cdc11 D65T compared to Cdc12 T75A or when the two mutations were combined (Figure 4B). As for nucleotide-exchanged conditions, bundling was substantially enhanced when septins were polymerized in solution (Figure 4D). The majority of Cdc11 D65T filaments had 4-5-fold greater intensity (or “bundling factor”) compared to the WT. A large population of Cdc12 T75A and the double mutant displayed a bundling factor of 1-2X compared to the wild type (WT, Figure 4E). Consistent with increased lateral interactions, filament flexibility decreased, with cdc11 D65T displaying the highest rigidity corresponding to the most bundled filaments (Figure 4F). Remarkably, the double mutant protein showed a less severe phenotype than Cdc11 D65T alone, possibly due increased compatibility of the mutant G-domain conformations (Weems *et al.*, 2014).

**Figure 4.**
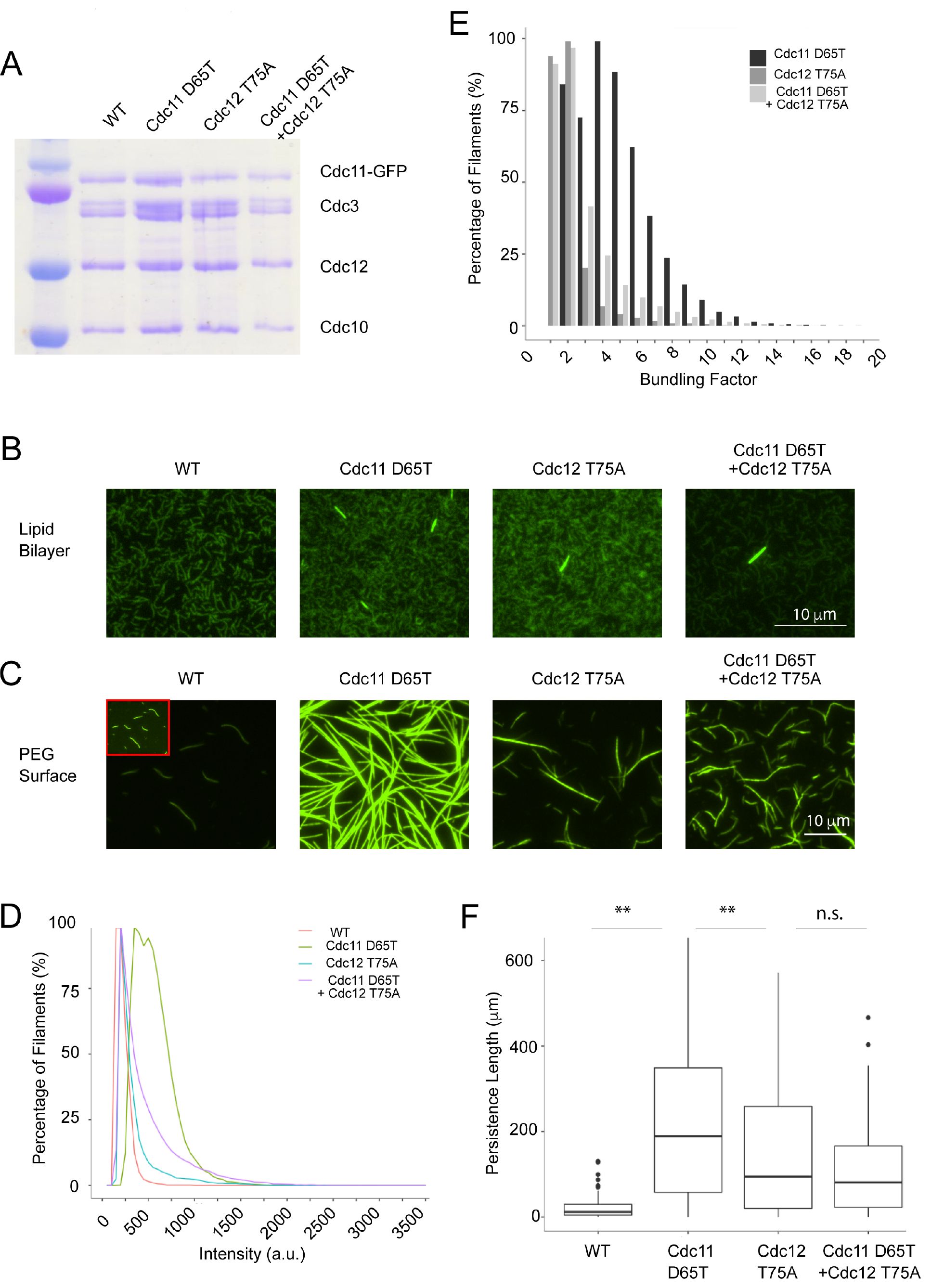
Nucleotide binding pocket mutants show an increase in bundling. (A) Coomassie-stained SDS gel showing no defects in complex formation by the mutants. (B) 5 nM protein was polymerized on a SLB in 50 mM salt. Mutants form bundled filaments. (C) 100nM septins were polymerized on a PEG surface in 50 mM salt. Mutants display increased lateral interactions in absence of a membrane. Insets show enhance contrasted filament. (D) Intensity distribution of filaments formed on PEG surface at 4°C. (E) Bundling factor of mutants polymerized on PEG compared to the WT. Percentage frequency captures filaments that showed at least a bundling of factor x. (F) Persistence length of WT and nucleotide mutant filaments (n ≥ 20 filaments, ** p<0.005, KS test). Mutants form rigid filaments compared to WT.

### Changing Shs1 proportion in branch collars results in polarity defects

A prediction of these *in vitro* experiments is that cells may be able to control features of higher-order assembly based on the relative composition of complexes of different identities and/or local nucleotide exchange. *A. gossypii* forms spatially-distinct septin structures that form at sites of different cellular curvatures (DeMay *et al.*, 2009; Bridges *et al.*, 2016) and, therefore, serves as a backdrop to examine how cells locally exploit different ratios of distinct complexes. Indeed, unlike in S. *cerevisiae*, Shs1 is essential for viability suggesting that modulating this terminal position is critical in these more morphologically complex cells (unpublished data). *A. gossypii* cells predominantly contain two types of higher-order septin structures: inter-region rings and
branch-associated collars. We found that although Cdc11 was generally more abundant than Shs1 in higher-order structures, the ratio of Shs1/Cdc11 appeared the same in both types of assemblies (Figure 5A).

**Figure 5.**
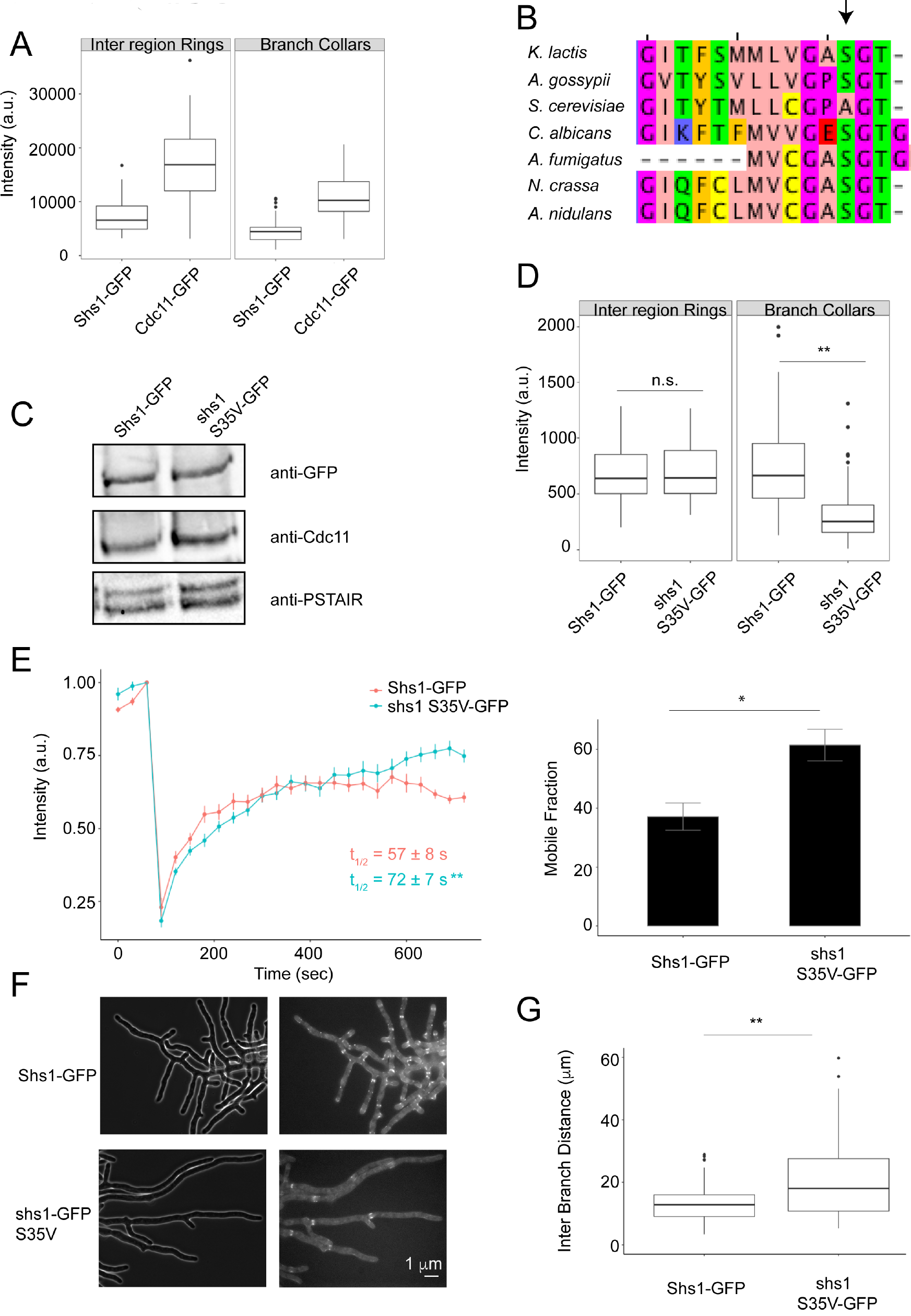
Analysis of Shs1:Cdc11 ratios and function of nucleotide state in *A. gossypii.* (A) Cdc11 and Shs1 intensities in different higher-order septin structures. (B) Sequence alignment showing conservation of serine residue predicted to be important for GTP binding/hydrolysis in different fungi (arrow) (C) Western blot of *A. gossypii* extract showing total Shs1 and Cdc11 levels. (D) GFP fluorescence intensity in inter-region rings (n ≥ 35) and branch collars (n ≥ 80) for Shs11-GFP and shs1 S35V-GFP. (E) *shs1 S35V-GFP* has slower dynamics but higher mobile fraction than Shs1-GFP at branch collars (n ≥ 8 rings). (E) Representative images of Shs1-GFP and *shs1 S35V-GFP.* (F) Inter branch distance in shs1-GFP and *shs1 S35V-GFP* cells (n ≥ 108 branches). * p < 0.05, ** p < 0.005, KS test.18

To examine the effect of nucleotide we mutated a serine residue in the GTP-binding site of Shs1 (*shs1 S35V*) that is conserved in other fungi but not in *S. cerevisiae* (Figure 5B). The mutant expression level was comparable to WT Shs1 and did not alter Cdc11 abundance (Figure 5C). Interestingly, we detected no inter-region ring intensity differences between *Shs1-GFP* and *shs1 S35V-GFP*, but observed a significant decrease in shs1 S35V-GFP incorporation at branch collars compared to Shs1-GFP (Figure 5D, p<0.005 KS test). FRAP experiments demonstrated that Shs1-GFP S35V recovered more slowly with a higher mobile fraction than Shs1-GFP (Figure 5E), suggesting oligomeric interactions and/or curved membrane affinity is perturbed. We also observed branch formation defects, revealed by increased inter-branch distance (Figure 5, F and G). Although the Cdc11:Shs1 ratio is comparable in different septin structures in WT cells, the nucleotide mutant selectively compromised Shs1 incorporation into branch collars and altered cell morphology.

We show that septin biophysical properties can be modulated by varying the incorporation of different types of complexes and the available nucleotide. Changing the Cdc11:Shs1 complex ratio can control filament length, flexibility, and dynamics and that nucleotide can influence filament bundling capacity. This work indicates multiple factors contribute to septin filament biophysical properties, and these help explain the diverse septin structures found inside cells.

## Materials and methods

### Protein expression and purification

BL21 (DE3) *E. coli* cells were transformed with duet septin expression plasmids immediately prior to production. Protein expression was induced by adding 1 mM IPTG at OD_600_ 0.6-0.7 at 22°C for 24 h with shaking. Cells were then harvested by centrifugation and pellets were either stored at −80°C until lysis or lysed immediately. Cells were incubated with lysis buffer (50 mM KH_2_PO_4_ pH 8.0, 1 M KCl, 1 mM MgCl_2_, 1% Tween-20, 10% glycerol, 1X protease inhibitor (Roche), and 20 mM imidazole) and 1 mg/ml lysozyme for 30 min on ice. Cells were then sonicated twice for 20 s each and the lysate was collected by spinning at 4°C for 30 min at 15,000 rpm using a SS-34 rotor in a Sorvall RC-6 centrifuge. The clarified lysate was filtered through a 0.45 μm filter before loading onto an equilibrated Ni^2+^-NTA agarose (Thermo Scientific) column. Bound protein was washed three times with wash buffer (50 mM KH_2_PO_4_ pH 8.0, 1 M KCl, and 20 mM imidazole) before elution by high imidazole buffer (50 mM KH_2_PO_4_,pH 8.0, 1 M KCl, and 500 mM imidazole). Protein was then dialyzed into septin storage buffer (50 mM Tris pH 8.0, 300 mM KCl and 1 mM β-mercaptoethanol) by using a 100,000 MW cutoff dialysis cassette (Thermo Fisher Scientific). Where applicable, the 6XHIS tag on Cdc12 was subsequently cleaved using ProTEV Plus Protease (Promega) and the protein was run over the Ni^2+−^NTA column to remove the cleaved 6XHIS tag. Protein was again dialyzed into septin storage buffer. Purity was analyzed by SDS gel electrophoresis and the total protein concentration was determined using Bradford reagent.

### Transmission Electron Microscopy

After purification, septin complexes were diluted to a concentration of 5 μg/ml in septin storage buffer. The diluted samples were adsorbed onto Formvar carbon coated grids (Ted Pella, Inc) after glow discharge. The grids were then washed twice with water and stained with 2% uranyl acetate.

Electron micrographs of the prepared samples were taken using an electron microscope (JEOL 1230 TEM) operated at 120 kV. Micrographs were acquired at 150,000X magnification using a CCD camera (SC1000; Gatan Orius). Length measurements were performed using Fiji (Schindelin *et al.*, 2012).

### Single molecule analysis using recombinant protein

Recombinantly-expressed proteins were diluted to pM concentrations before being immobilized on a clean glass surface. Glass coverslips were prepared by sequential sonication steps of 10 mins each in 3 N KOH, 100% ethanol, and ddH_2_O. The coverslips were washed with ddH_2_O five times between each step. Coverslips were then dried and plasma cleaned for 15 min. Plastic chambers were glued onto coverslips. Septins were diluted in buffer containing 50 mM Tris pH 8, 300 mM KCl, 1 mg/ml BSA and 1mM β-mercaptoethanol. The chamber was washed once with buffer containing 50 mM Tris pH 8, 300 mM KCl, 1 mg/ml BSA and 1 mM β-mercaptoethanol before adding septins to a final pM concentration in the chamber. The protein was allowed to settle for 5 min before imaging.

### Single Molecule Pull-down (SiMPull) on yeast extract

SiMPull was performed as previously described (Jain *et al.*, 2009). Yeast extracts were prepared by growing 1 L of *S. cerevisiae* cells to an OD of 0.6-0.7 in YPD medium with selection. Cells were then spun down at 5000 x g for 15 min and pellets were re-suspended in 1 ml of YPD. The yeast cells were then flash frozen in liquid nitrogen and ground using a coffee grinder. The ground paste was suspended in lysis buffer containing 50 mM Hepes pH 7.6, 1 mM KCl, 1 mM EGTA, 5% glycerol, 0.01% Tween and 1X protease inhibitor mix. Bradford reagent was used to determine protein concentration of the lysates. A goat α-rabbit biotin conjugated (Invitrogen) secondary antibody was immobilized on biotin PEG coated coverslips. Rabbit α-GFP (Invitrogen) was used to recruit Cdc11-GFP from the extracts. Single molecules were imaged using time lapse acquisition with an exposure time and frame rate of 200 ms.

### Supported Lipid Bilayer (SLB) preparation

Glass coverslips were cleaned and SLB’s were prepared with 75% (mol%) 1,2-dioleoyl-sn-glycero-3-phosphocholine (DOPC; Avanti Polar Lipids 850375) and 25% (mol%) L-α-phosphatidylinositol (liver, bovine; Avanti Lipids 840042) as described previously (Bridges *et al.*, 2014). Proteins were added to a final concentration of 5 nM total protein in Reaction Buffer A containing 50 mM Tris pH 8.0, 50 mM KCl, 1 mg/ml fatty acid free BSA (Sigma A6003) and 1 mM β-mercaptoethanol

### Septin polymerization on PEG surface

To investigate septin polymerization in the absence of a lipid bilayer, purified septins were allowed to polymerize on PEG-passivated coverslips. Plastic chambers were glued onto the coverslip and septins were added to a final concentration of 100 nM in Reaction Buffer B containing 50 mM Tris pH 8, 50 mM KCl, 1 mg/ml fatty acid free BSA, 0.1% methyl cellulose and 1mM β-mercaptoethanol. Septins were allowed to polymerize for ~ 1 h before imaging unless otherwise specified.

### Septin polymerization in presence of Mg^2+^ and/or nucleotide

To determine the effect of magnesium on septin polymerization, 5 nM total protein was allowed to polymerize in Reaction Buffer A + 1 mM MgCl_2_ on a SLB. For polymerization on PEG, 100 nM total protein was polymerized in Reaction Buffer B + 3 mM MgCl_2_.

Nucleotide exchange for septins was done by adopting previously published methods (Sheffield *et al.*, 2003; Huang *et al.*, 2006). 250 nM total septin protein was incubated with 250 μM nucleotide and 1 mM MgCl_2_ in high salt (300 mM type of salt?) for 3 h at 30°C to allow nucleotide exchange. For SLB experiments, the protein was diluted to a final concentration of 5 nM total protein, 5 μM nucleotide and 20 μM MgCl_2_ in Reaction Buffer A and allowed to polymerize for ~ 30 min before imaging.

For polymerization on a PEG surface, higher protein concentration was typically required to obtain filaments that can be visualized using TIRF due to 3-D instead of 2-D diffusion. For this reason, the same exchange protocol was applied but using a higher starting protein concentration. 400 nM total protein was incubated with 400 μM nucleotide and 3 mM MgCl_2_ in high salt for 3 hours at 30°C. For polymerization, the protein was diluted to a final concentration of 100 nM, 100 μM nucleotide and 1 mM MgCl_2_ in Reaction Buffer B. The protein was allowed to polymerize overnight on PEG before imaging.

### Total Internal Reflection Fluorescence Microscopy

Septin filaments on supported lipid bilayers, PEG passivated glass, and single molecules were imaged using TIRF microscopy to minimize background. Images were acquired on a Nikon TiE TIRF system using an Apo 100X/1.49 NA oil objective equipped with a solid state laser system (15 mW; Nikon LUn-4) and a sCMOS camera (Photometrics Prime 95B).

### Preparing SLB microspheres and curvature analysis

To measure septin adsorption on different curved surfaces, SLB-covered microspheres were prepared as described previously (Bridges *et al.*, 2016). Small unilamellar vesicles (SUVs) were prepared with a lipid composition of 75% 1,2-dioleoyl-sn-glycero-3-phosphocholine (DOPC; Avanti Polar Lipids 850375), 25% L-α-phosphatidylinositol (liver, bovine; Avanti Lipids 840042) and >0.1% L-α-phosphatidylethanolamine-N-(lissamine rhodamine B sulfonyl) (Rh-PE; egg-transphosphatidylated, chicken; Avanti Polar Lipids 810146). SUVs were then adsorbed onto silica microspheres (0.96, 3.17 and 5.06 μm; Bangs Laboratories). Plastic chambers were glued onto PEG-passivated coverslips. Septins were then incubated with the lipid coated bead mixture at a final concentration of 100 nM (unless otherwise stated) for 1 h at room temperature to reach equilibrium, in buffer containing 50 mM Tris pH 8, 100 mM KCl, 1 mg/ml fatty acid free BSA, 0.1% methyl cellulose and 1 mM β-mercaptoethanol. Images were acquired using a Nikon TiE widefield with a Plan Apo 100X/1.45 NA oil objective equipped with a sCMOS camera (Andor Zyla 4.2 plus). Septin binding analysis was done in Imaris 8.1.2 (Bitplane) by creating surfaces on the beads and determining the sum intensity for both the lipid and the septin channel. Septin adsorption was calculated by dividing septin sum intensity by the lipid septin intensity. Subsequent analysis and plots were generated using RStudio version 1.0.136. Statistical analysis was done by performing a Kruskal-Wallis rank sum test followed by a Bonferonni-adjusted Dunn test as previously described (Bridges *et al.*, 2016).

### Cdc11 and Shs1 ratio experiments

Different proportions of purified Cdc11 and Shs1 complexes were diluted in septin storage buffer before addition to the SLB at a final protein concentration of 5 nM. Septin polymerization was followed for a total of 15 min using TIRF (described above). Short videos to determine persistence length were acquired with a time interval of 250 ms and a 100 ms exposure.

For curvature analysis using different Cdc11 and Shs1 proportions, the two complexes were mixed in varying proportion in septin storage buffer to a total protein concentration of 400 nM. For curvature binding the mixtures containing different proportions of Cdc11 and Shs1 were added to the beads to a total protein concentration of 400 nM. The rest of the experiment was performed as described above for curvature analysis.

### Fragmentation, annealing, and bundling frequency analysis

For fragmentation, annealing, bundling, and unbundling analysis, isolated filaments were identified and followed for ~50 frames and all of these events were recorded. Each interaction was considered authentic and not transient if it persisted at least 2 frames. If a filament changed length considerably after an annealing event, it was given a new identity for tracking purposes. Initial filament lengths were recorded. For each filament tracked the number of events was divided by the starting length of the filament to obtain a measure of annealing, fragmentation, bundling, or unbundling per unit length.

### Bundling factor computation

For bundling analysis, “tophat” morphological filtering was applied to the images before thresholding to create filament masks in MATLAB R2016a (MathWorks). Using the masks each isolated filament was identified and a frequency distribution of intensity was determined. To account for noise, intensity values which accounted for less than 2% of the filament size were not considered. The 2% threshold was empirically selected as a strong criterion for rejection of noise-based outliers. The intensity ranges were then divided by the average intensity of the WT or untreated filaments to acquire the distribution of the metric we term “bundling factor.”

### Single particle tracking

Particle centers were segmented within time-lapse data using a convolutional neural network (Newby *et al.*, 2017). Linking particles across consecutive frames and matching particles between channels was formulated as a bipartite graph matching problem. Linking particles across consecutive frames and matching particles between channels was formulated as a bipartite graph matching problem, a common optimization problem that selects the best match between two sets of points that minimizes the total cost, in our case distance. The matching problem was solved using the Hungarian-Munkres algorithm with a cutoff distance of five pixels (Newby *et al.*, 2017), i.e. particles were required to be closer than five pixels apart in consecutive frames to be a valid match. Maximum particle intensities were plotted for display.

### Persistence length analysis

Filament persistence length was determined as described by Bridges et al. 2014, using previously published methodology for actin and microtubules (Gittes, 1993). 15 or more filaments were selected with little change in length and tracked for ≥ 6 frames. 2-D coordinates over the length of the filament were determined using Fiji. MATLAB was then used to determine and report the median persistence length of the filaments. Plots were generated using the RStudio ggplot2 package.

### Western Blotting

A. *gossypii* cultures were grown in *Ashbya* Full Medium (AFM) containing antibiotics and selection for 15-17 h at 30°C with shaking. Cells were collected by vacuum filtration, suspended in lysis buffer (50 mM Hepes pH 7.8, 1 M KCl, 1 mM MgCl_2_, 1 mM EGTA, 0.1% Tween, 2X protease inhibitor, 5% glycerol) and lysed using a bead beater. Lysate was collected by spinning at 13,000 rpm for 10 min at 4°C. Bradford assay was used to determine the lysate protein concentration. Equal amounts of protein were loaded onto a SDS gel for western blotting. The membranes were probed with rabbit polyclonal α-GFP (Invitrogen) for Shs1-GFP and shs1 S35V-GFP, rabbit polyclonal α-Cdc11 (Santa Cruz) for Cdc11 and rabbit α-PSTAIR (Abcam) as a loading control, all at 1:1000 dilution.

### FRAP microscope setup and analysis

A. *gossypii* was grown as above. FRAP experiments were performed on a Nikon Eclipse Ti with a Yokogawa spinning disc using a Plan Apo 100X/1.49 NA oil objective. Photobleaching of septin branch collars was performed using a 405 nm laser (450 mW) in conjunction with a mosaic micro-mirror array system. Images were acquired using an EM-CCD camera (Hamamatsu). Individual branch collars for each strain were bleached and imaged every 30 s for 12 min and the fluorescence intensity was recorded at each time point.

To measure septin branch collar intensity over time, a region of interest (ROI) was drawn around the ring and the mean fluorescence was measured. The mean fluorescence of an adjacent area in the cytosol was determined and subtracted from the septin intensity measurements to account for background. ROI position was adjusted to account for branch collars moving due to cell growth. There was minimum photo-bleaching over the course of the experiment. The mobile fractions were determined by determining the ratio of percent recovery to percent bleach. t_1/2_ was calculated using the formula

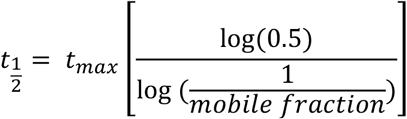

where t_max_ is the time at which recovery plateaus. Plots were generated using the ggplot2 package in RStudio (Wickham et al., 2009).

### *A. gossypii* growth and imaging

A. *gossypii* cultures were grown in Ashbya full medium (AFM) containing antibiotics and appropriate selection for 15-17 hours at 30°C with shaking. To image GFP-tagged septins, cells were collected by centrifugation at low speed and washed twice with low-fluorescence minimal media and mounted onto 2% agarose gel pads made of low-fluorescence minimal media. Images were acquired on a Zeiss Axioimage-M1 upright light microscope (Carl Zeiss, Jena, Germany) equipped with a Plan-Apochromat 63X/1.4 NA oil objective and an Exfo X-Cite 120 lamp. Z-stacks were acquired using a step size of 0.3 μm and ~20 slices per image. Image analysis was performed in Fiji. Septin structure intensity was determined by drawing a ROI and taking the sum intensity measurements from 5 different points along the structure. An adjacent area of the cytosol was measured using the same ROI for 5 different regions and the mean sum intensity of the cytosol was subtracted from the mean sum intensity of the septin structures to account for background subtraction. Plots were generated using the ggplot2 package in RStudio.

### Strain Construction

Strains, plasmids, and oligonucleotides used in this study are listed in Table S1. Custom oligonucleotides were from Integrated DNA Technologies (IDT, Coralville USA), and Phusion polymerase was used for PCR (NEB, Ipswitch USA). To generate plasmid pMVB128 HIS-Sccdc12 T75A/ScCDC10 (AGB561), AGO1340 and AGO1341 were used to introduce the point mutation using the Agilent XL QuickChange kit. This created an additional EcoRV site at 349 bp. The correct product was verified using digestion with EcoRV and sequencing using AGO1192 and AGO1342. To generate plasmid pMVB133 ScCDC3/Sccdc11 D65T-GFP (AGB562), primers AGO1347 and AGO1348 were used to introduce the point mutation as well as a SalI site at 4004 bp using the Agilent XL QuickChange kit. The correct product was verified by digestion with SalI and HindIII as well as sequencing using primers AGO1349 and AGO1350.

To generate plasmid pMVB133 ScCDC3/ScSHS1-GFP (AGB548), assembly cloning was used. ScSHS11 was PCR amplified using genomic DNA and primers AGO1273 and AGO1274 to produce the insert fragment. The 1712 bp product was gel extracted using the E.Z.N.A gel extraction kit (Omega Bio-Tek, Norcross USA). The plasmid vector was generated by PCR using AGB454 and primers AGO1271 and AGO1272, to generate a 6511 bp product. The PCR product was purified by gel extraction. The vector and the insert were ligated using NEB HiFi DNA Assembly Master Mix. The resulting plasmid was isolated from transformants using the EZNA Plasmid Mini Kit (Omega Bio-Tek). The correct product was verified by digestion with HindIII and SacI. To generate plasmid pCOLA ScSHS1-GFP (AGB895) assembly cloning was used. The insert was generated by PCR using AGB598 as template and primers AGO1807 and AGO1808 to get a 944 bp product and was treated with DpnI to digest the template. The vector was generated through PCR using AGB560 as template and primers AGO1805 and AGO1806 to produce a 5282 bp product. Vector was purified by gel extraction. The vector and insert were assembled using NEB HiFi DNA Assembly Master Mix. The resulting plasmid was isolated by mini kit and tested by NruI and PacI double digestion. The correctly digested plasmid was sequence verified using AGO1496 and AGO1320.

To generate AGshs1 S35V-GFP strain (AG372.1), primers AGO537 and AGO538 were used to introduce the S35V point mutation in plasmid AGB88 using the Stratagene QuickChange II XL site directed mutagenesis kit (AGB211). AGO541 and AGO412 were used to verify the point mutation. The plasmid AGB211 was linearized by cutting with BlpI and the 5263-nucleotide fragment was transformed into AG416. Genomic DNA was isolated using a Qiagen DNeasy Plant Mini kit (Qiagen, Venlo Netherlands). Heterokaryons and homokaryons were verified using AGO592 and AGO593.

### Sequence Alignment

Protein sequences were BLASTED against *Ashbya gossypii* Shs1 using BLASTp on the Saccharomyces Genome Database. The protein that showed the highest consensus with the BLAST query amongst the fungal species listed in Figure 5 were selected. Sequence alignment was performed using ClustalW ((Thompson *et al.*, 2002; Goujon *et al.*, 2010). JalView (Version 2.10.2b2) was used to display and view the sequence alignment (Waterhouse *et al.*, 2009).

## Acknowledgements

We thank all Gladfelter lab members, J. Moseley, E. Smith and H. Higgs for useful discussions; V Madden for support with TEM work; M Baig for help with bundling analysis; and S. Dundon for critically reading the manuscript.

This work was supported by the National Science Foundation (MCB MCB-0719126).

The authors declare no competing financial interests.

## Supplementary materials

**Table S1:**
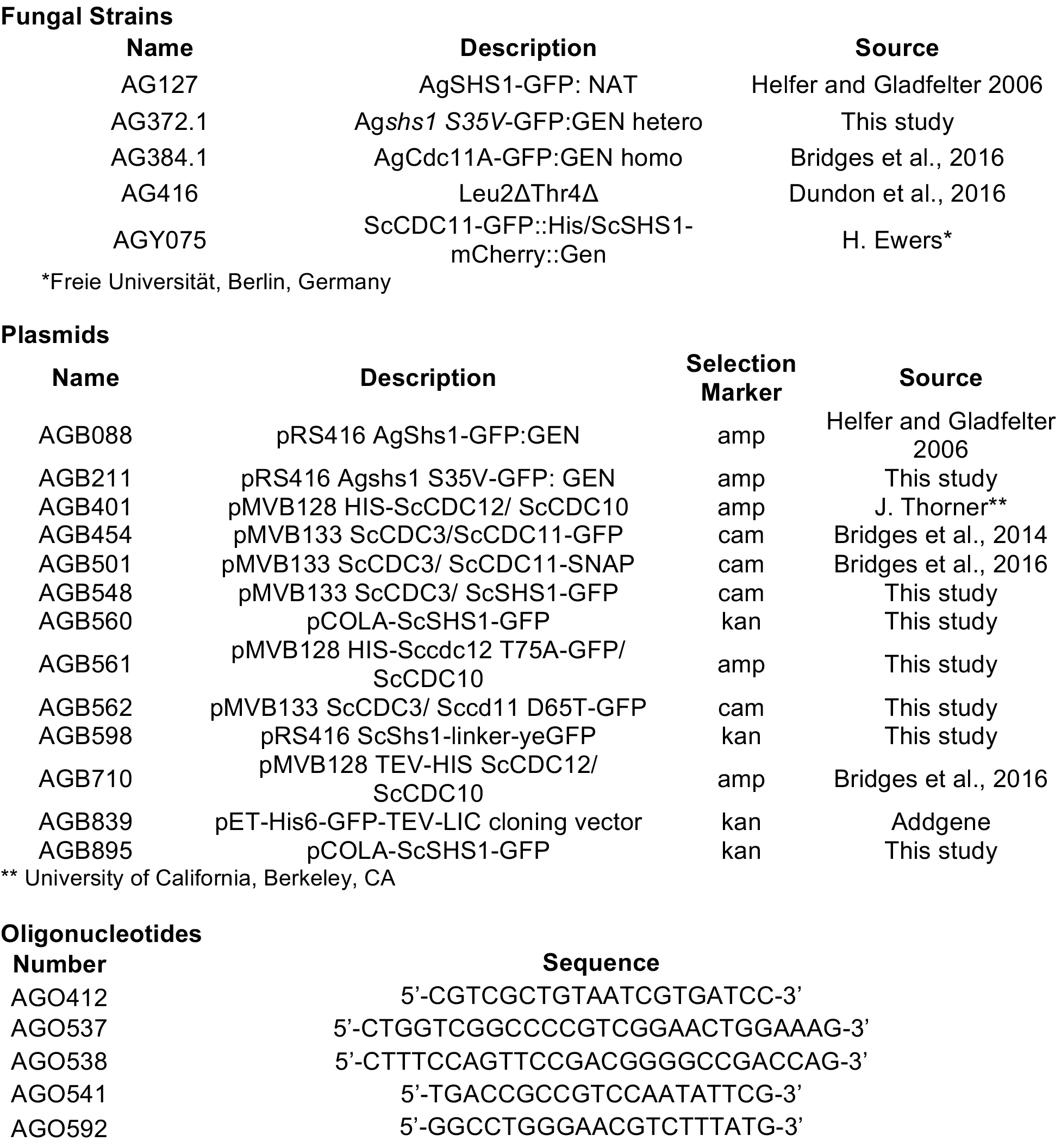
A list of all the fungal strains, plasmids and oligonucleotides that were used in the study.

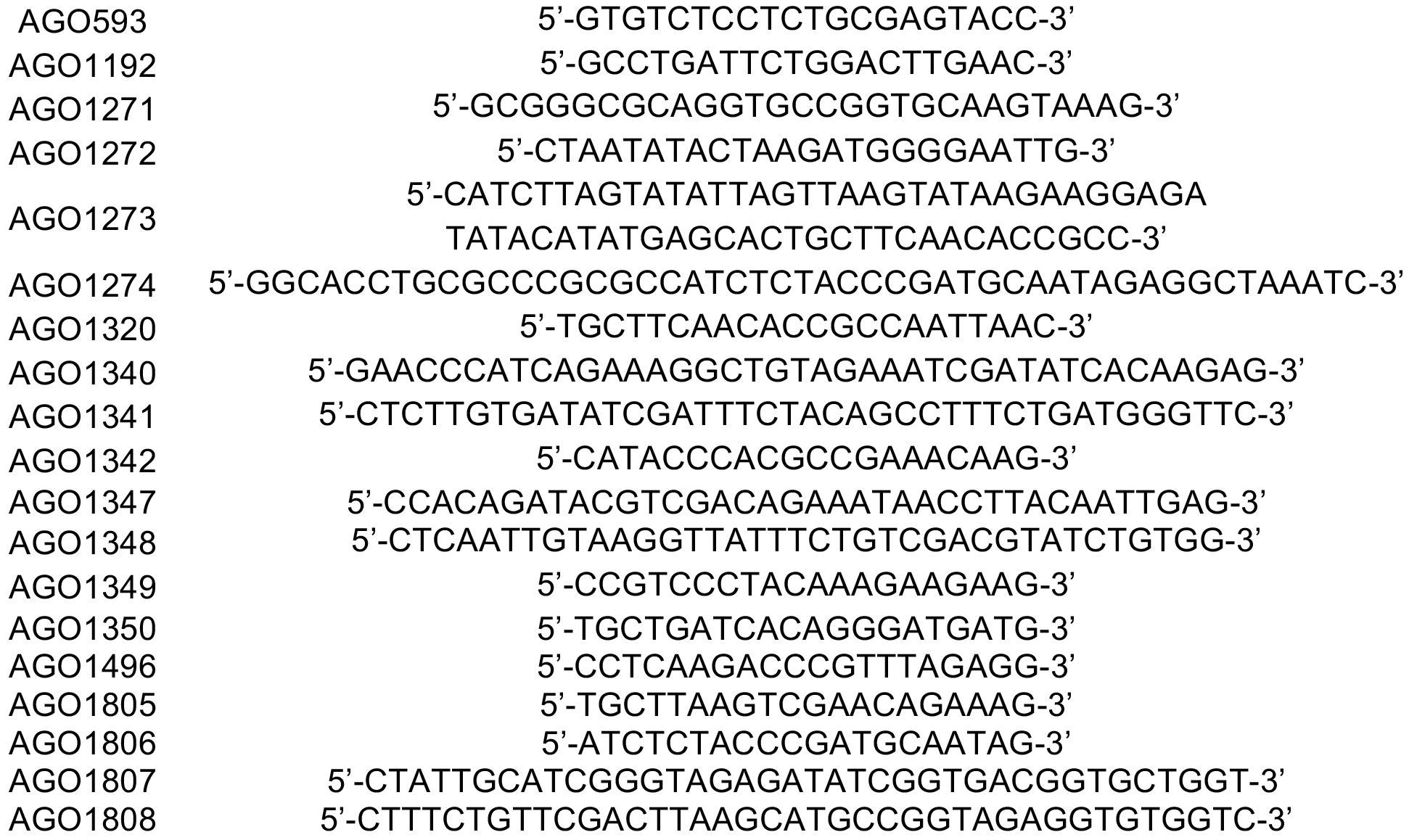

Supplementary Movie- Video 1- Filaments formed on a SLB by addition of 5 nM total protein containing Cdc11 only complexes, corresponding to Figure 2. Time lapse was acquired via TIRF microscopy with a 100 ms exposure and a 250 ms interval.

Supplementary Movie- Video 2- Filaments formed on a SLB by mixing Cdc11 and Shs1 complexes in a 1:1 ratio to a final protein concentration of 5 nM, corresponding to Figure 2. Video was acquired with a 100 ms exposure and 250 ms time interval.

Supplementary Movie- Video 3- Filaments formed in absence or presence of nucleotide using 5 nM protein and 5 μM nucleotide, corresponding to Figure 3.

## References

Akhmetova, K., Balasov, M., Huijbregts, R. P. H., and Chesnokov, I. (2015). Functional insight into the role of Orc6 in septin complex filament formation in Drosophila. Mol. Biol. Cell 26, 15–28.

Alvarez-Tabarés, I., and Pérez-Martín, J. (2010). Septins from the phytopathogenic fungus Ustilago maydis are required for proper morphogenesis but dispensable for virulence. PLoS ONE 5, e12933–17.

Baumann, S., König, J., Koepke, J., and Feldbrügge, M. (2014). Endosomal transport of septin mRNA and protein indicates local translation on endosomes and is required for correct septin filamentation. EMBO Rep 15, 94–102.

Bertin, A., McMurray, M. A., Thai, L., Garcia, G., III, Votin, V., Grob, P., Allyn, T., Thorner, J., and Nogales, E. (2010). Phosphatidylinositol-4,5-bisphosphate promotes budding yeast septin filament assembly and organization. Journal of Molecular Biology 404, 711–731.

Booth, E. A., and Thorner, J. (2016). A FRET-based method for monitoring septin polymerization and binding of septin-associated proteins. In: Septins, Elsevier, 35–56.

Booth, E. A., Vane, E. W., Dovala, D., and Thorner, J. (2015). A Förster Resonance Energy Transfer (FRET)-based System Provides Insight into the Ordered Assembly of Yeast Septin Hetero-octamers. J. Biol. Chem. 290, 28388–28401.

Bridges, A. A., Jentzsch, M. S., Oakes, P. W., Occhipinti, P., and Gladfelter, A. S. (2016). Micron-scale plasma membrane curvature is recognized by the septin cytoskeleton. The Journal of Cell Biology 213, 23–32.

Bridges, A. A., Zhang, H., Mehta, S. B., Occhipinti, P., Tani, T., and Gladfelter, A. S. (2014). Septin assemblies form by diffusion-driven annealing on membranes. Proc. Natl. Acad. Sci. U.S.A. 111, 2146–2151.

Chong, Y. T., Koh, J. L. Y., Friesen, H., Duffy, S. K., Cox, M. J., Moses, A., Moffat, J., Boone, C., and Andrews, B. J. (2015). Yeast proteome dynamics from single cell Imaging and automated analysis. Cell 161, 1413–1424.

DeMay, B. S., Meseroll, R. A., Occhipinti, P., and Gladfelter, A. S. (2009). Regulation of distinct septin rings in a single cell by Elm1p and Gin4p kinases. Mol. Biol. Cell 20, 2311–2326.

Field, C. M. (1996). A purified Drosophila septin complex forms filaments and exhibits GTPase activity. The Journal of Cell Biology 133, 605–616.

Finnigan, G. C., Takagi, J., Cho, C., and Thorner, J. (2015). Comprehensive genetic analysis of paralogous terminal septin subunits Shs1 and Cdc11 in Saccharomyces cerevisiae. Genetics 200, 821–841.

Ford, S. K., and Pringle, J. R. (1991). Cellular morphogenesis in the Saccharomyces cerevisiae cell cycle: localization of the CDC11 gene product and the timing of events at the budding site. Dev. Genet. 12, 281–292.

Garcia, G., III, Bertin, A., Li, Z., Song, Y., McMurray, M. A., Thorner, J., and Nogales, E. (2011). Subunit-dependent modulation of septin assembly: Budding yeast septin Shs1 promotes ring and gauze formation. The Journal of Cell Biology 195, 993–1004.

Gittes, F. (1993). Flexural rigidity of microtubules and actin filaments measured from thermal fluctuations in shape. The Journal of Cell Biology 120, 923–934.

Goujon, M., McWilliam, H., Li, W., Valentin, F., Squizzato, S., Paern, J., and Lopez, R. (2010). A new bioinformatics analysis tools framework at EMBL-EBI. Nucleic Acids Res. 38, W695–W699.

Hernández-Rodráguez, Y., Masuo, S., Johnson, D., Orlando, R., Smith, A., Couto-Rodriguez, M., and Momany, M. (2014). Distinct septin heteropolymers co-exist during multicellular development in the filamentous fungus Aspergillus nidulans. PLoS ONE 9, e92819–10.

Huang, Y. -W., Surka, M. C., Reynaud, D., Pace-Asciak, C., and Trimble, W. S. (2006). GTP binding and hydrolysis kinetics of human septin 2. Febs J. 273, 3248–3260.

Jain, A., Ramani, B., Raghunathan, K., Jena, P., Xiang, Y., and Ha, T. (2009). Single Molecule Immuno Pull Down Assay (SiMPull) for Studying Protein-Protein Interactions. Biophysical Journal 96, 25a.

Kaplan, C. et al. (2015). Absolute Arrangement of Subunits in Cytoskeletal Septin Filaments in Cells Measured by Fluorescence Microscopy. Nano Letters 15, 3859–3864.

Kim, M. S., Froese, C. D., Estey, M. P., and Trimble, W. S. (2011). SEPT9 occupies the terminal positions in septin octamers and mediates polymerization-dependent functions in abscission. The Journal of Cell Biology 195, 815–826.

Marrington, R., Small, E., Rodger, A., Dafforn, T. R., and Addinall, S. G. (2004). FtsZ fiber bundling Is triggered by a conformational change in bound GTP. J. Biol. Chem. 279, 48821–48829.

Nagaraj, S., Rajendran, A., Jackson, C. E., and Longtine, M. S. (2008). Role of nucleotide binding in septin-septin interactions and septin localization in *Saccharomyces cerevisiae*. Molecular and Cellular Biology 28, 5120–5137.

Newby, J. M., Schaefer, A. M., Lee, P. T., Forest, M. G., and Lai, S. K. (2017). Deep neural networks automate detection for tracking of submicron scale particles in 2D and 3D. 1–14.

Schindelin, J. et al. (2012). Fiji: an open-source platform for biological-image analysis. Nature Methods 9, 676–682.

Sellin, M. E., Sandblad, L., Stenmark, S., and Gullberg, M. (2011). Deciphering the rules governing assembly order of mammalian septin complexes. Mol. Biol. Cell 22, 3152–3164.

Sheffield, P. J., Oliver, C. J., Kremer, B. E., Sheng, S., Shao, Z., and Macara, I. G. (2003). Borg/Septin interactions and the assembly of mammalian septin heterodimers, trimers, and filaments. J. Biol. Chem. 278, 3483–3488.

Sirajuddin, M., Farkasovsky, M., Zent, E., and Wittinghofer, A. (2009). GTP-induced conformational changes in septins and implications for function. Proc. Natl. Acad. Sci. U.S.A. 106, 16592–16597.

Tang, J. X., and Janmey, P. A. (1996). The polyelectrolyte nature of F-actin and the mechanism of actin bundle formation. J. Biol. Chem. 271, 8556–8563.

Thompson, J. D., Gibson, T. J., and Higgins, D. G. (2002). Multiple sequence alignment using ClustalW and ClustalX, Hoboken, NJ, USA: John Wiley & Sons, Inc.

Versele, M., Gullbrand, B., Shulewitz, M. J., Cid, V. J., Bahmanyar, S., Chen, R. E., Barth, P., Alber, T., and Thorner, J. (2004). Protein-protein interactions governing septin heteropentamer assembly and septin filament organization in Saccharomyces cerevisiae. Mol. Biol. Cell 15, 4568–4583.

Vrabioiu, A. M., Gerber, S. A., Gygi, S. P., Field, C. M., and Mitchison,T. J. (2004). The majority of the Saccharomyces cerevisiae septin complexes do not exchange guanine nucleotides. J. Biol. Chem. 279, 3111–3118.

Waterhouse, A. M., Procter, J.B., Martin, D. M. A., Clamp, M., and Barton, G. J. (2009). Jalview Version 2–– a multiple sequence alignment editor and analysis workbench. Bioinformatics *25*, 1189–1191.

Weems, A. D., Johnson, C. R., Argueso, J.L., and McMurray, M. A. (2014). Higher-order septin assembly is driven by GTP-promoted conformational changes: evidence from unbiased mutational analysis in Saccharomyces cerevisiae. Genetics 196, 711–727.

Weems, A., and McMurray, M. (2017). The step-wise pathway of septin hetero-octamer assembly in budding yeast. eLife 6, 20007.

Zander, S., Baumann, S., Weidtkamp-Peters, S., and Feldbrügge, M. (2016). Endosomal assembly and transport of heteromeric septin complexes promote septin cytoskeleton formation. J. Cell. Sci. 129, 2778–2792.

